## Supplementary Material for "Inhibition of the serine protease HtrA1 by SerpinE2 suggests an extracellular proteolytic pathway in the control of neural crest migration"

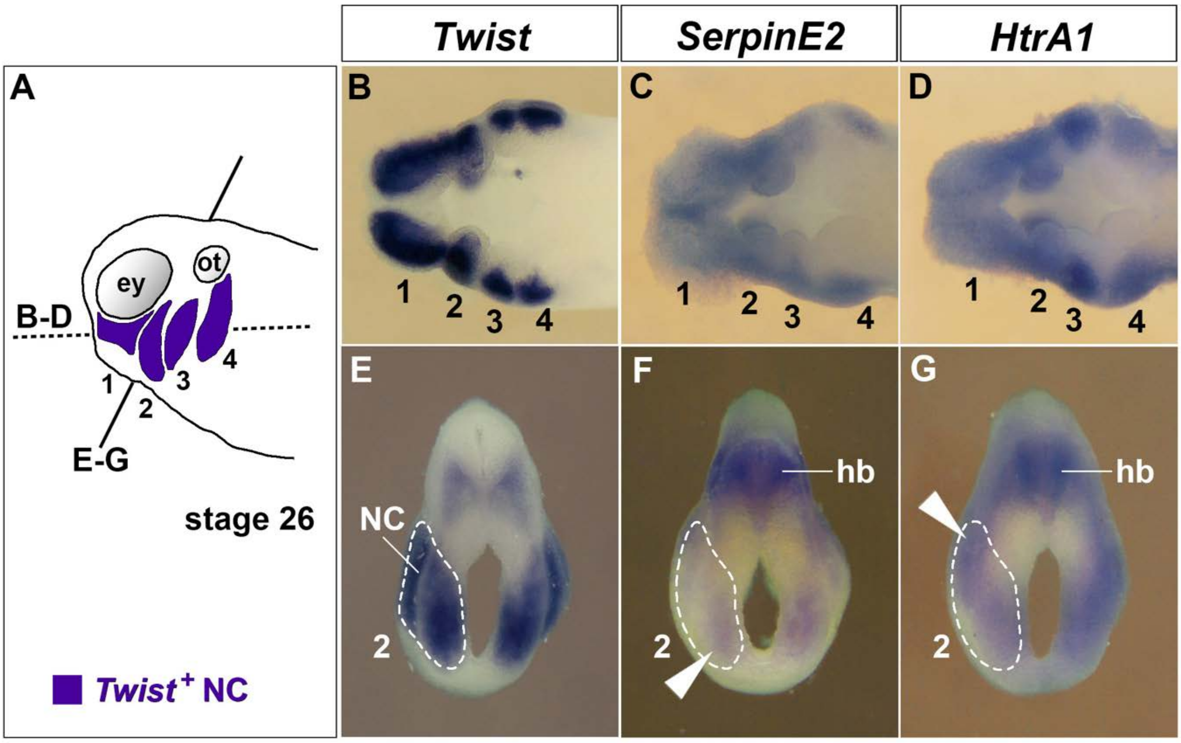


**Figure 1-figure supplement 1. *Twist*, *SerpinE2* and *HtrA1* gene expression in sections of tailbud stage embryos after whole-mount *in situ* hybridization**

**(A)** Schematic of embryo at stage 26. Labelled in violet are the *Twist*-positive NC cells in the mandibular arch (1), hyoid arch (2), anterior branchial arch (3), and posterior branchial arch (4). Indicated are the level of horizontal sections in panels B-D (strippled line) and the level of transversal sections in in panels E-G (continuous line).

**(B-D)** Horizontal hemisections viewed from dorsal.

**(E-G)** Transversal sections. The strippeled line labels the *Twist*$^{+}$ NC. Note the arrowheads indicating the ventral *SerpinE2* and the dorsal *HtrA1* expression domains in the NC of the hyoid arch.

ey, eye; hb, hindbrain; ot, otic placode.

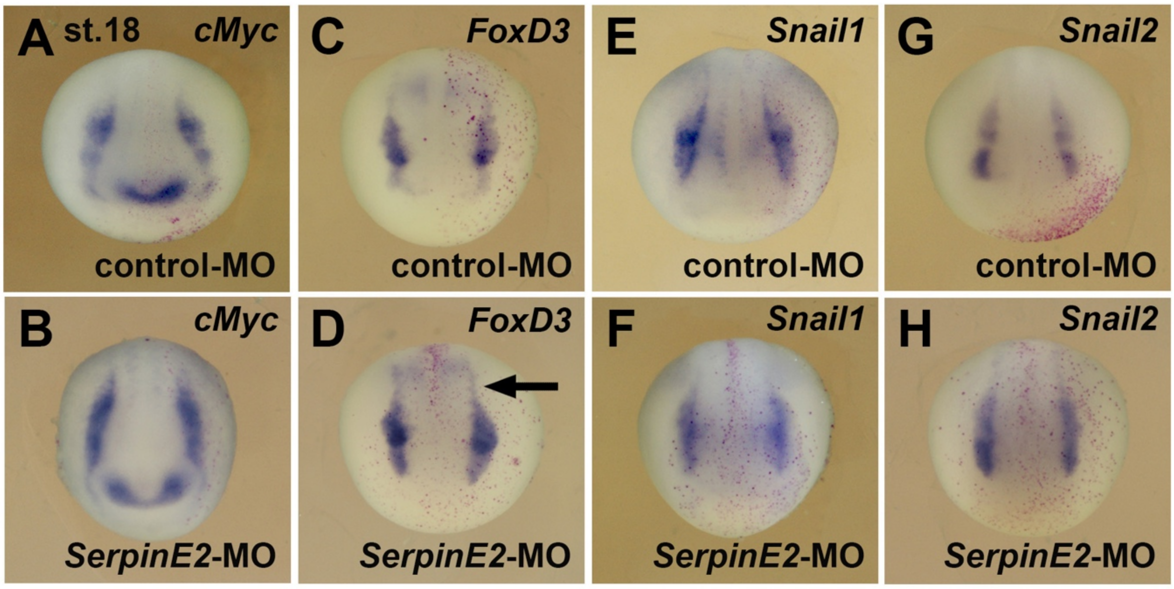


**Figure 2-figure supplement 1. SerpinE2 depletion does not affect the specification of neural crest cells**

**(A-H)** Anterior view of embryos at stage 18 after whole-mount *in situ* hybridization. Note that neither control-MO (A,C,E,G) nor *SerpinE2*-MO (B,D,F,H) affect the expression of the neural crest cell markers *cMyc*, *FoxD3*, *Snail1*, and *Snail2*. The arrow demarcates the trunk neural crest.

Embryos were injected into a single animal blastomere at the 8-cell stage with 10 ng morpholino oligonucleotide (MO) and 100 pg *nlacZ* mRNA as lineage tracer (red nuclei). Indicated phenotypes were shown in A, 5/5; B, 9/9; C, 8/8; D, 9/9; E, 6/6; F, 6/7; G, 8/8; H, 5/6.

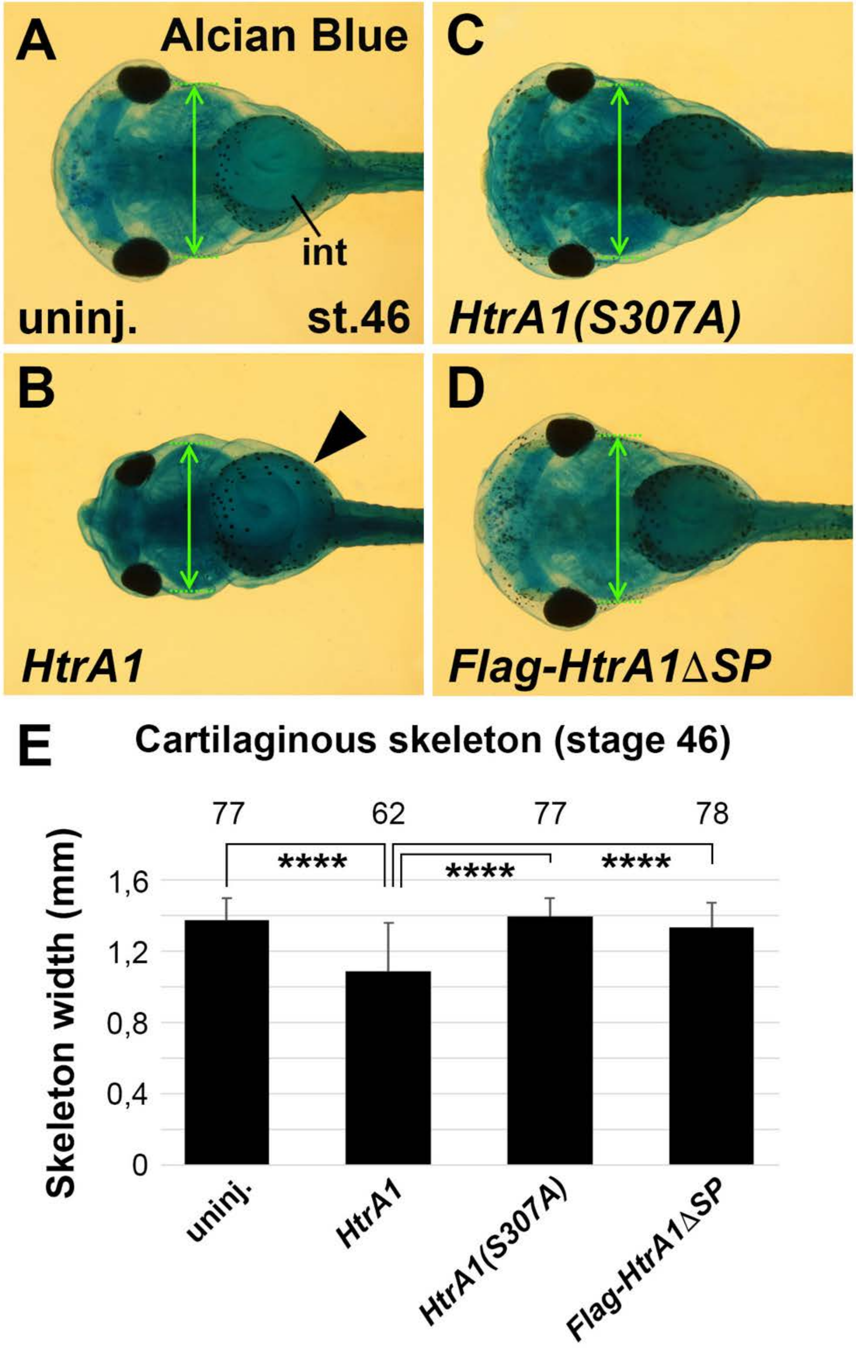


**Figure 4-figure supplement 1. Overexpression of HtrA1 decreases cartilaginous elements and craniofacial structures.**

Embryos were injected into four animal blastomeres at the 8-cell stage with a total of 100 pg mRNA.

**(A-D)** Ventral view of embryos at stage 46 after Alcian Blue staining. Note that *HtrA1* mRNA decreases the skeleton width (double arrow at the level of the ceratobranchial cartilage structures) but does not affect gut coiling (arrowhead). *HtrA1(S307A)* and *HtrA1ΔSP* mRNAs have no effects.

**(E)** Quantification of the skeleton width. The number of analyzed specimen is indicated above each column.

int, intestine. Indicated phenotypes were shown in A, 73/74; B, 58/62; C, 73/77; D, 67/78 embryos. At least two experiments were done.

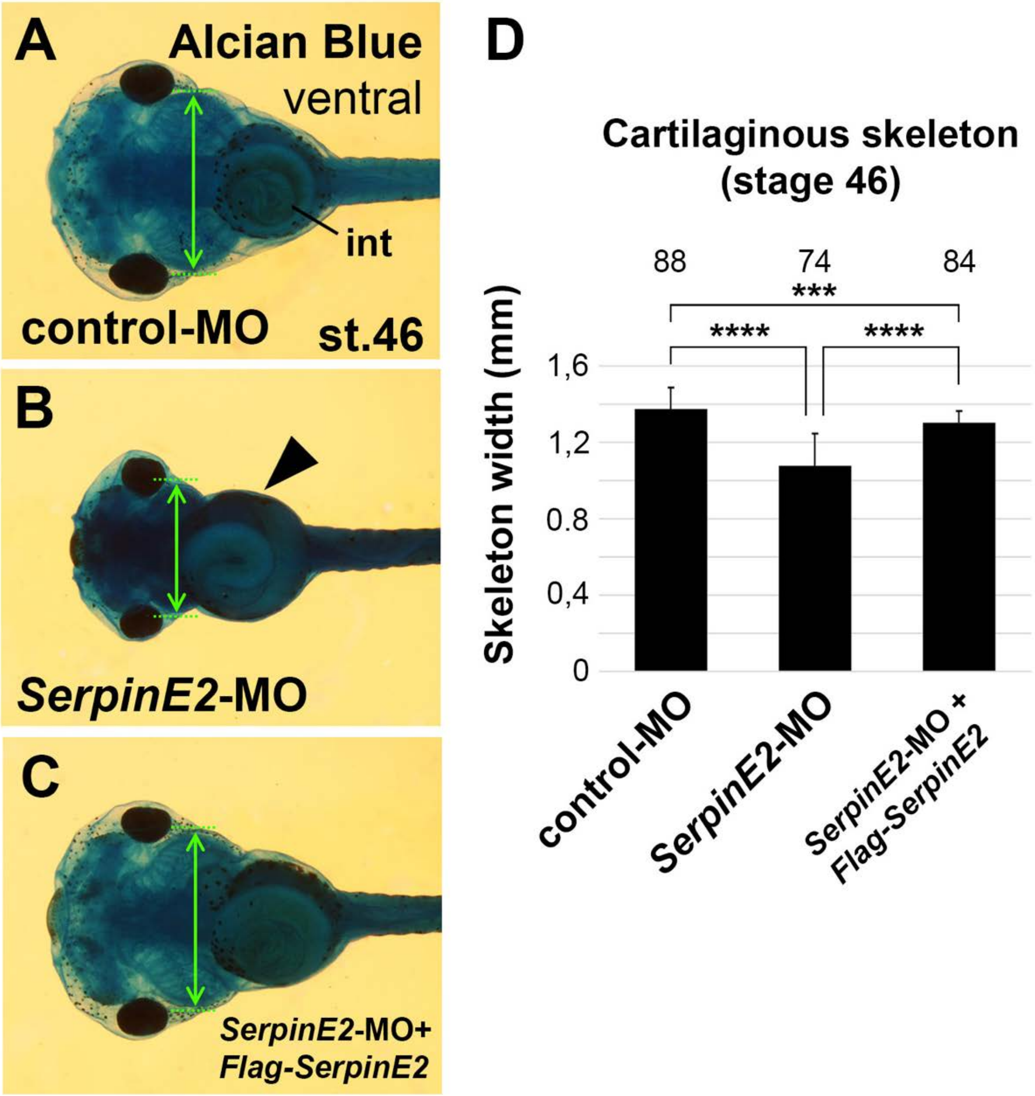


**Figure 4-figure supplement 2. Knockdown of SerpinE2 reduces craniofacial skeleton formation.**

Morpholino oligonucleotides (MOs) were injected into *Xenopus* embryos at the 8-cell stage. **(A-C)** Ventral view of tadpoles at stage 46 after Alcian Blue staining. Note that *SerpinE2*-MO decreases the branchial skeleton width (double arrow) but does not affect gut coiling (arrowhead). The control-MO and a combination of *SerpinE2*-MO and *Flag-SerpinE2* mRNA have no effects.

**(D)** Quantification of the skeleton width. The number of analyzed specimen is indicated above each column. Embryos were injected into four animal blastomeres with a total of 40 ng MO.

cb, ceratobranchial; ch, ceratohyal; et, ethmoid-trabecular; int, intestine; mc, Meckel's cartilage; pq, palatoquadrate. Indicated phenotypes were shown in A, 76/88; B, 72/74; C, 78/85. At least two experiments were done. At least three experiments were done.

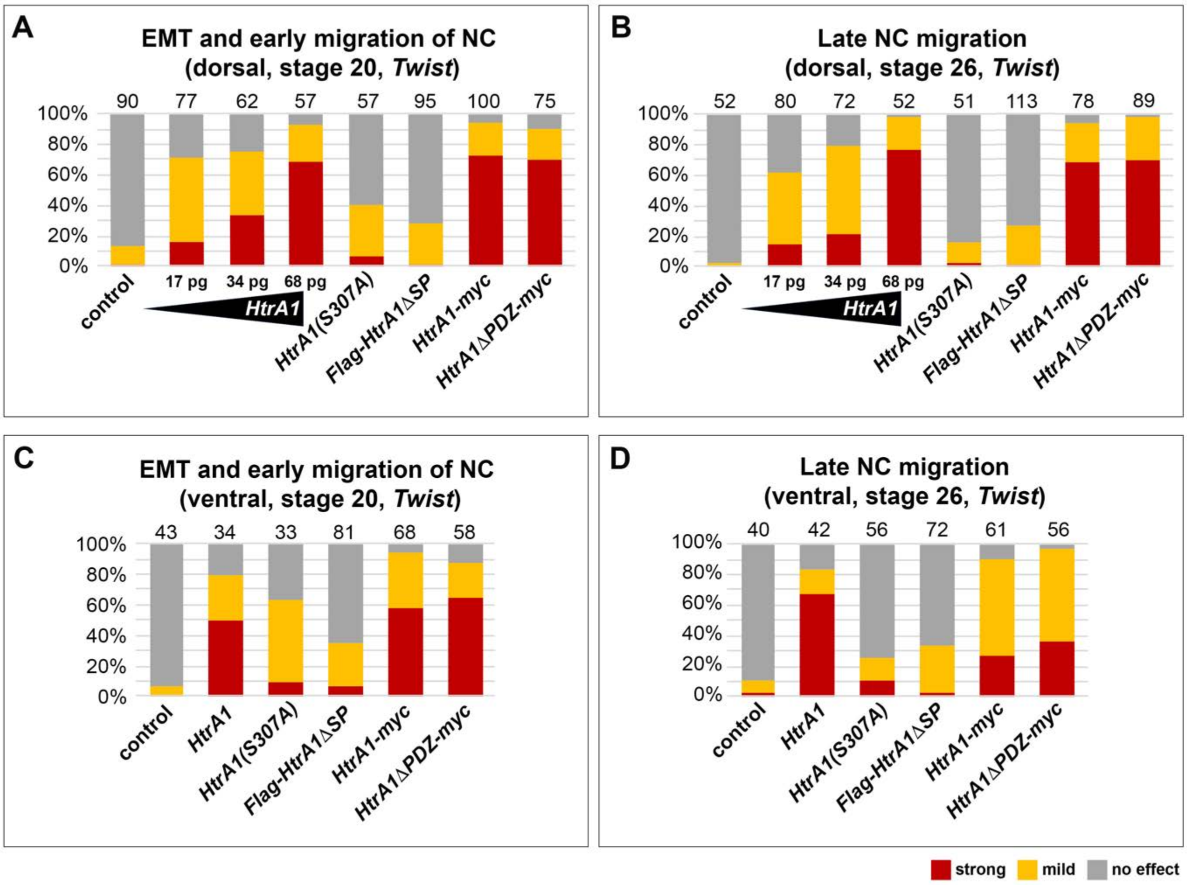


**Figure 6-figure supplement 1. *HtrA1* mRNA inhibits neural crest migration in a concentration-dependent manner.**

If not otherwise indicated, embryos were injected with 65 pg *HtrA1*-derived mRNAs into a single dorsal animal blastomere at the 8-cell stage. Quantification of EMT and migration of *Twist*$^{+}$ NC cells in embryos at stages 20 and 26. Defects were assessed based on a comparison of injected embryos and non-injected control siblings. Normal phenotypes are distinguished from phenotypes with either mild or strong defects in NC cell migration. A mild defect is defined as a reduction by more than 25% in at least one NC segment. A strong defect is a reduction by more than 50% in all NC segments. The number of analyzed embryos per sample is indicated above the columns. At least two experiments were done.

**(A,B)** Following dorsal injection, *HtrA1* mRNA blocks EMT and cell migration of NC cells in a dose-dependent manner. Note that *HtrA1-myc* and *HtrA1∆PDZ-myc* mRNAs cause migration defects with a similar strength, whereas HtrA1(S307A) and *Flag-HtrA1∆SP* mRNAs have only little effects.

**(C,D)** Ventral injection of *HtrA1* mRNA cause relatively mild NC migration defects.

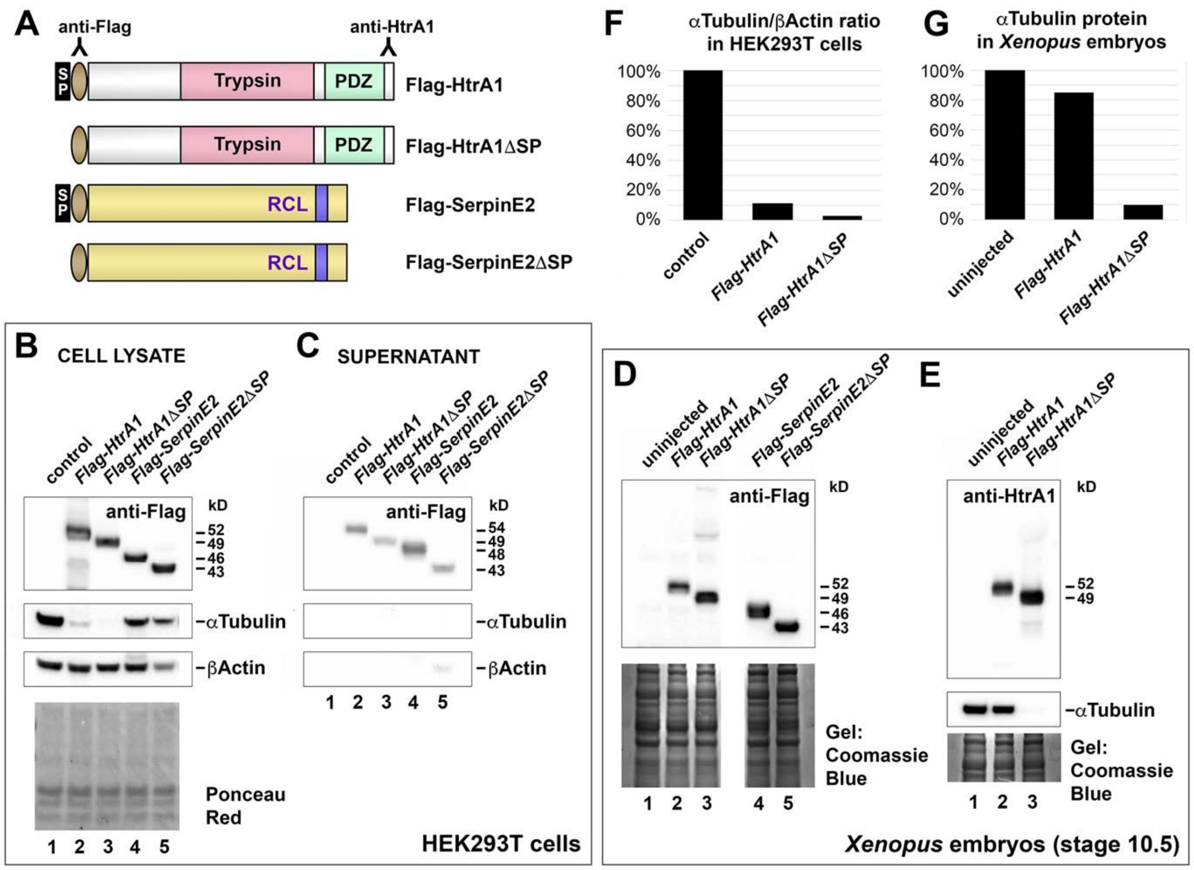


**Figure 6-figure supplement 2. Cytoplasmic HtrA1 causes reduction of αTubulin protein levels in mammalian cells and *Xenopus* embryos.**

**(A)** Schematic representation of the protein constructs. Indicated are antibody binding sites. RCL, reactive center loop; SP, signal peptide.

**(B,C)** Western blot analysis of cell lysate and culture supernatants. HEK293T cells were transiently transfected with pCS2 vector constructs containing Flag epitope-tagged cDNAs encoding HtrA1, HtrA1ΔSP, SerpinE2, and SerpinE2ΔSP. Note that Flag-HtrA1 and Flag-SerpinE2 have higher molecular weights in the supernatant (54 and 48 kD) than in the cell lysate (52 and 46 kD), indicating that these proteins are glycosylated in the secretory pathway. Flag-HtrA1ΔSP and Flag-SerpinE2ΔSP are less abundant in the supernatant, and their molecular weights (49 and 43 kD) the same as in the cell lysate, suggesting that these signal peptide-deficient proteins are not secreted and their low levels in the supernatant likely due to cell lysis. Empty vector was used as negative control. Ponceau Red staining of membrane shows equal amounts of protein in the lysate samples.

**(D,E)** Western blot analysis of *Xenopus* embryos at stage 10.5. Embryos were injected into the animal pole at the 4-8 cell stage. Injected mRNA doses per embryo were 320 pg (*HtrA1*-derived) and 4 ng (*SerpinE2*-derived). Coomassie blue staining shows equal protein loading in the gels.

**(F,G)** Quantification of αTubulin protein levels relative to βActin in HEK293T cells (F) and relative to Ponceau Red staining in *Xenopus* embryos at stage 10.5 (G). Experiments were done at least twice.

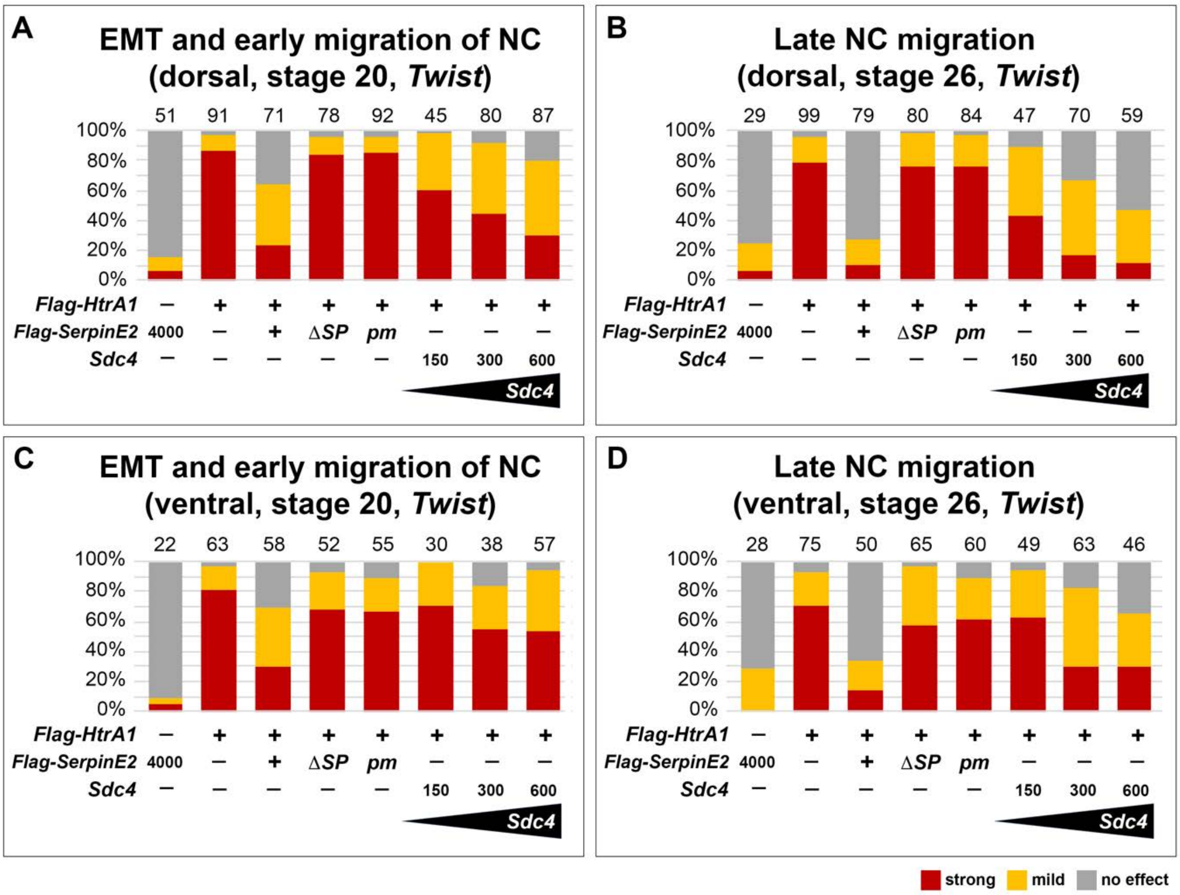


**Figure 7-figure supplement 1. *SerpinE2* and *Sdc4* mRNAs partially rescue neural crest migration defects that are induced by HtrA1 overexpression.**

Quantification of EMT and migration of *Twist*$^{+}$ NC cells in embryos at stages 20 and 26. Defects were assessed based on a comparison of injected embryos and non-injected control siblings. The number of analyzed embryos per sample is indicated above the columns. Note that secreted Flag-SerpinE2, but not the cytosolic Flag-SerpinE2ΔSP and mutant Flag-SerpinE2pm constructs, revert NC migration defects in *Flag-HtrA1* mRNA-injected embryos.

**(A,B)** Dorsal injection of *Sdc4* mRNA rescues *Flag-HtrA1*-induced migration defects in a dose-dependent manner.

**(C,D)** *Sdc4* mRNA is less efficient in rescuing NC migration upon ventral co-injection with *Flag-HtrA1* mRNA.

Embryos were injected into one animal blastomere at the 8-cell stage with mRNA doses of 65 pg (*Flag-HtrA1*) and 200 pg (*SerpinE2*-derived). Indicated *Sdc4* mRNA amounts per embryo are in pg. Experiments were done at least three times.

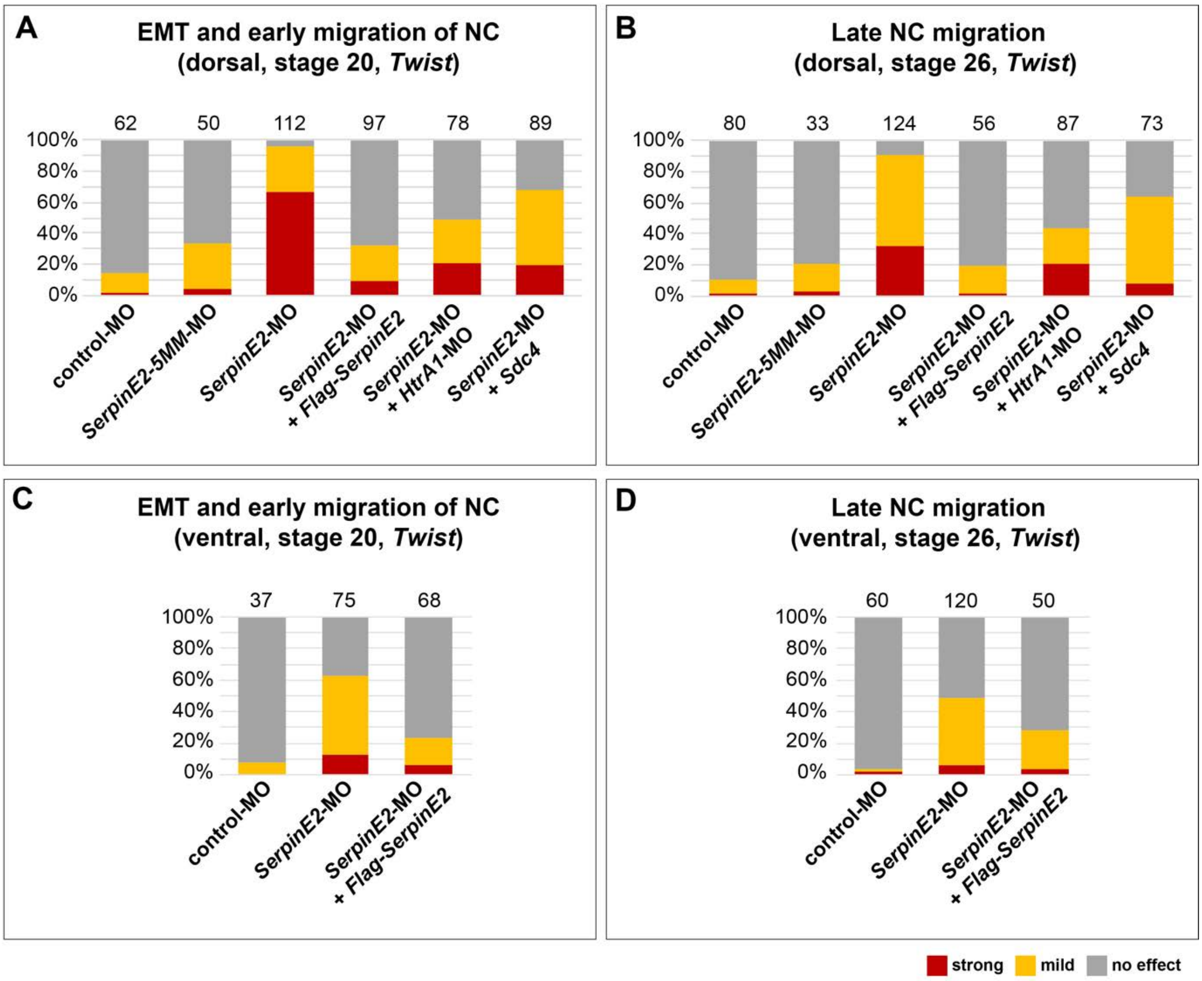


**Figure 8-figure supplement 1. SerpinE2-MO causes neural crest migration defects which are rescued by *HtrA1*-MO and *Sdc4* mRNA.**

EMT and NC migration defects were quantified in embryos at stages 22 and 26 after whole-mount in situ hybridization with the NC marker *Twist*. Defects were assessed based on a comparison between the injected and non-injected side within the same specimen. Normal phenotypes were distinguished from phenotypes with either mild or strong defects in NC cell migration. A mild defect is defined as a reduction by more than 25% in at least one NC segment. A strong defect is a reduction by more than 50% in all NC segments. The number of analyzed embryos per sample is indicated above the columns.

**(A, B)** Dorsal injection of *SerpinE2*-MO blocks EMT and migration of NC cells, while control-MO and *SerpinE2*-5MM-MO have no or only little effect. *Flag-SerpinE2* mRNA, *HtrA1*-MO, and *Sdc4* mRNA restore normal NC migration in *SerpinE2*-morphant embryos.

**(C,D)** Upon ventral injection, *SerpinE2*-MO mildly blocks EMT and migration of NC cells. Morpholino oligonucleotides (MOs, 10 ng) and mRNAs were injected into one animal blastomere at the 8-cell stage. Injected mRNA doses per embryos were 333 pg (*Flag-SerpinE2*) and 300 pg (*Sdc4*). Experiments were done at least three times.

**Videos**

**Figure 5-video 1. Collective migration of neural crest cells *in vitro*.** Embryos were injected into the animal pole at the 4-cell stage with 40 ng standard control-MO. The cranial NC was explanted at stage 16/17, plated on fibronectin and imaged 7 hours after culture. Filming was done for 20 minutes. Note that cells at the leader front of the cluster extend lamellipodia and filopodia.

**Figure 5-video 2. Knockdown of SerpinE2 prevents adhesion and migration of neural crest cells.** Embryos were injected into the animal pole at the 4-cell stage with 40 ng *SerpinE2*-MO. The cranial NC was explanted at stage 16/17, plated on fibronectin and imaged 7 hours after culture. Filming was done for 10 minutes. Note that cells acquire a ball-like shape and freely float around.

**Source data files**

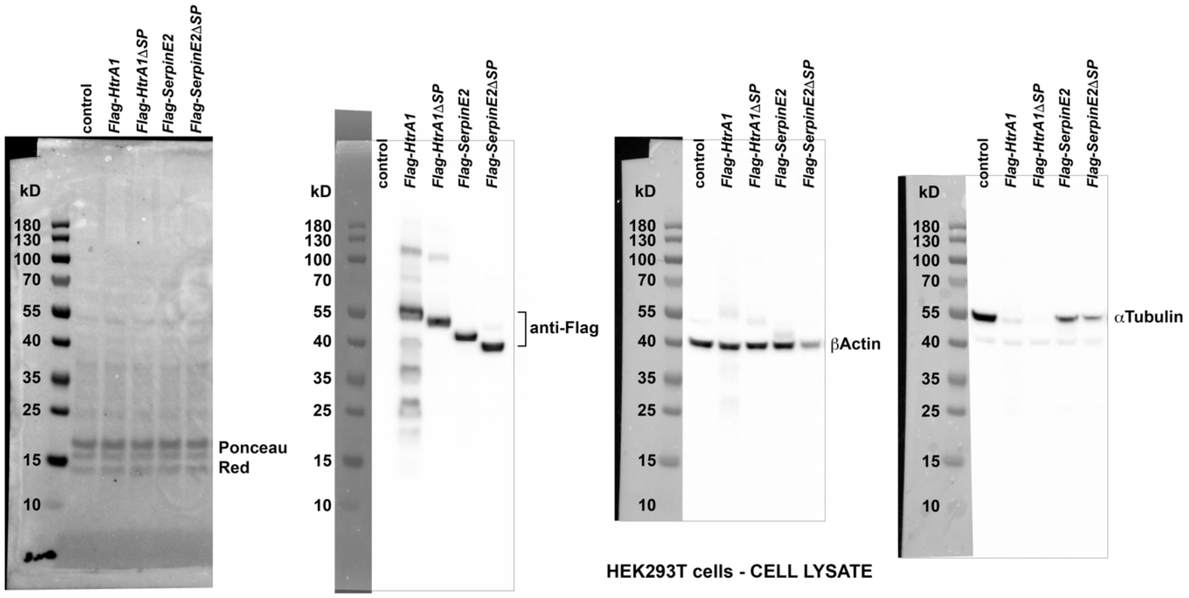


**Figure 6-Figure supplement 2-Source data 1. Western blot analysis of HEK293T cell lysate.** HEK293T cells were transfected with pCS2 vector alone as a control and pCS2 constructs containing Flag epitope-tagged cDNAs encoding HtrA1, HtrA1ΔSP, SerpinE2, and SerpinE2ΔSP. Ponceau Red staining validates that in all samples equal amounts of protein were transferred to the membrane. The same membrane was then sequentially incubated with antibodies against Flag epitope, βActin and αTubulin. Note that traces of anti-Flag antibody are seen in the βActin blot, and traces of βActin antibody in the αTubulin blot due to incomplete membrane stripping between the antibody incubations.

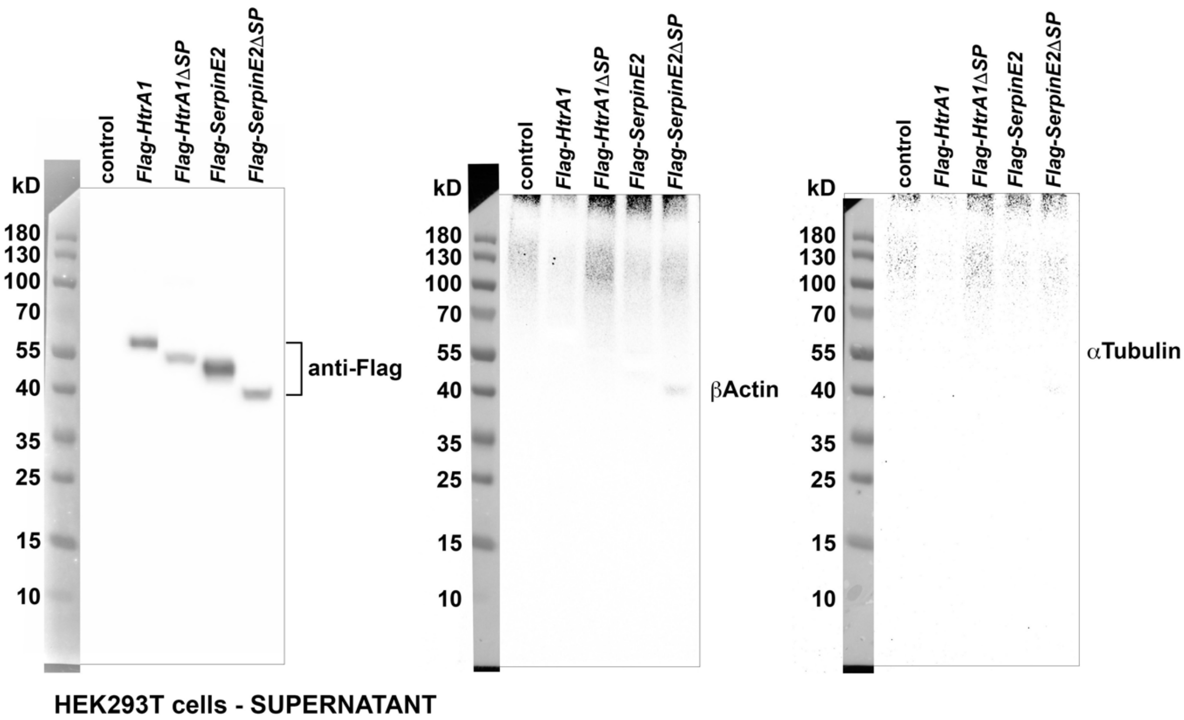


**Figure 6-Figure supplement 2-Source data 2. Western blot analysis of HEK293T cell supernatant.** The same membrane was sequentially incubated with antibodies against Flag epitope, βActin and αTubulin.

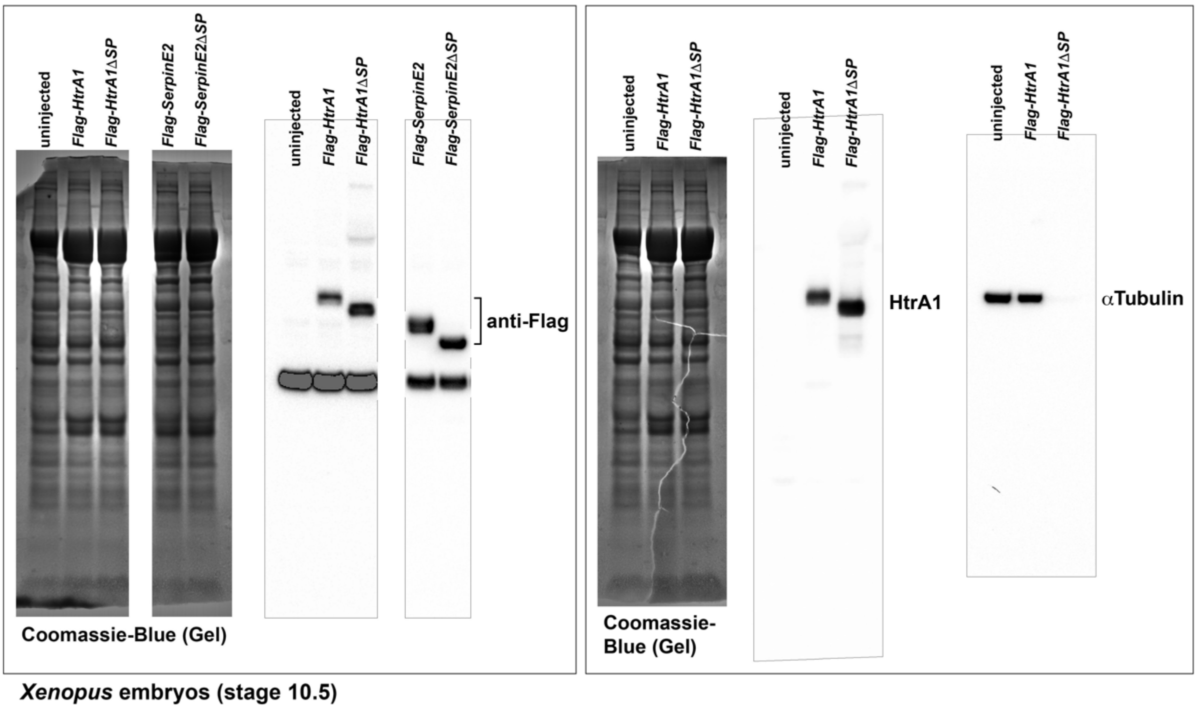


**Figure 6-Figure supplement 2-Source data 3. Western blot analysis of *Xenopus* embryos at stage 10.5.** *Xenopus* embryos were uninjected or microinjected with mRNA encoding Flag-tagged HtrA1, HtrA1ΔSP, SerpinE2, and SerpinE2ΔSP. Coomassie-Blue staining of the gel validates that in all samples equal amounts of protein were loaded. Membranes were incubated with anti-Flag antibody (left side) or sequentially with antibodies against HtrA1 and αTubulin (right side).
